# Delivery of collagen-encoding mRNA to the skin for cosmetic applications

**DOI:** 10.1101/2025.11.02.686162

**Authors:** Jianyu Wen, Bo Pang, Jinyu Han, Shuai Wang, Lianqiang Xu, Yufeng Wu, Hao Zhang, Fengyan Luo, Zelong Jin, Yong Hu

**Author notes:** These authors have contributed equally to this work (Jianyu Wen, Bo Pang).

## Abstract

Collagens, the predominant structural proteins in mammalian, maintain tissue integrity through its unique triple-helical conformation. Traditionally, cosmetic collagens have been administered as hydrolyzed peptides derived from animal sources, which exhibit poor bioavailability and functional efficacy due to limited skin penetration and disrupted tertiary/quaternary structures. To overcome these limitations, we developed a novel system for *in vivo* collagen expression. This system utilizes the topical application of *in vitro* synthesized, AI-designed mRNAs encoding types I, III, and XVII collagens to improve human skin condition. Significant upregulation of target collagen expression was observed in dermal fibroblasts, leading to improved skin condition through in situ assembly of collagens in natural conformation.

## 1 Introduction

Collagens comprise a family of at least 28 members, each containing a characteristic triple-helical domain [1]. As the most abundant extracellular proteins in mammals, collagens mediate cell-matrix interactions and form the structural scaffold of connective tissues such as skin, tendons, and ligaments[2]. Structurally, collagens are among the most complex natural polymers. The unique collagen protein consists of three left-handed polypeptide α chains that form a triple helix via hydrogen bonding, which further assembles into a right-handed superhelix through winding around themselves and their axis [1, 3-5].

In the skin, collagens occur naturally in the dermis as fibers, accounting for approximately 70% of the skin’s dry weight, and provide density and elasticity. Types I and III collagens are the major structural components of the skin, conferring mechanical strength, structural support, and elasticity. Type XVII collagen, a membrane-associated protein expressed in hemidesmosomes, regulates epidermal cell proliferation and is associated with skin aging when deficient [1, 6-8].

In cosmetic industry, collagens are widely used ingredients to improve the skin appearance [6, 9]. Conventional cosmetic collagen products are derived from animal sources in the form of hydrolyzed peptides, which suffer from poor bioavailability and functional performance due to inadequate skin penetration and loss of higher-order structures [10, 11]. To overcome these limitations, we developed a novel mRNA-based system that enables the *in vivo* expression and subsequent in situ assembly of collagens into their natural conformation.

## 2. Results

### 2.1 AI-based Sequence Optimization of mRNA

The transdermal delivery, stability and translation efficiency of mRNA not only depend on cis-elements such as 5’-UTR, 3’-UTR and poly(A) tail, but also depend on global sequence features such as GC content and secondary structure. Using an AI-driven platform by minimizing mRNA folding energy and ribosomal stalling, and by rational designing of optimal secondary structure to enhance delivery through intact human skin, we optimized mRNA sequences including CDS encoding collagens and cis-elements combinations in fibroblast cell HDF-α. The screening results for mRNA encoding collagen III were shown in Fig. 1. Compared with wild-type collagen III mRNA, the expression level of optimized CDS for collogen III were elevated significantly (Fig. 1A). To further improve expression efficiency in fibroblast cells, cis-elements combinations were screened based on AI-driven prediction (Fig. 1B). Collogen type III can be efficiently expressed in fibroblast cell line HDF-α when transfected with optimized mRNA (Fig. 2). Similar optimization processes were conducted for mRNA encoding collogen types I and XVII (The expression data in fibroblast cells shown in Supplemental Fig. 1). The effect of global sequence optimization on the transdermal delivery efficiency were evaluated and confirmed with *in vitro* test method of collagen I contents with fibroblasts according to Group Standard T/SHRH 031-2020 by Shanghai Daily Chemistry Trade Association[12] and T-Skin™, a reconstructed full thickness skin model[13] (data shown in Supplemental Fig. 2)

**Figure 1.**
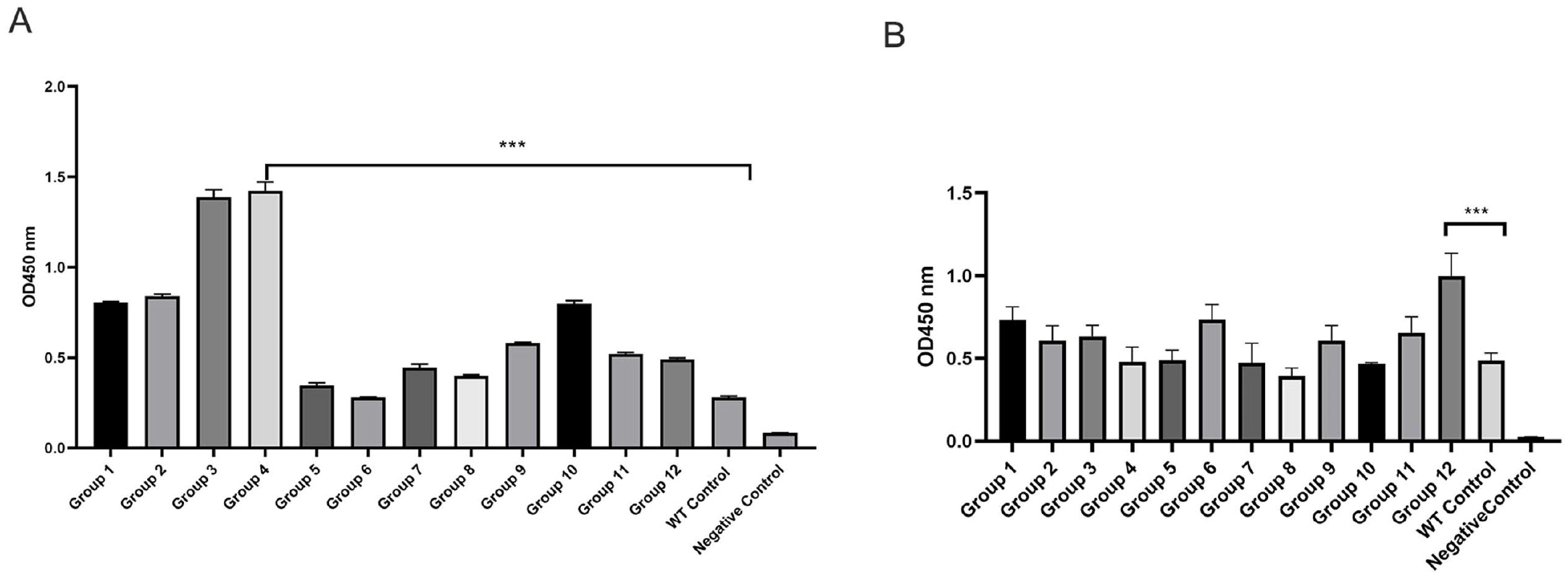
AI-assisted sequence design in fibroblast expression system. A Strategy screening for CDS optimization. B Screening of 5’UTR and 3’UTR regions. Data represent mean ± SD from three biological replicates (Student’s t test, *P < 0.05, **P < 0.01, ***P < 0.001).

**Figure 2.**
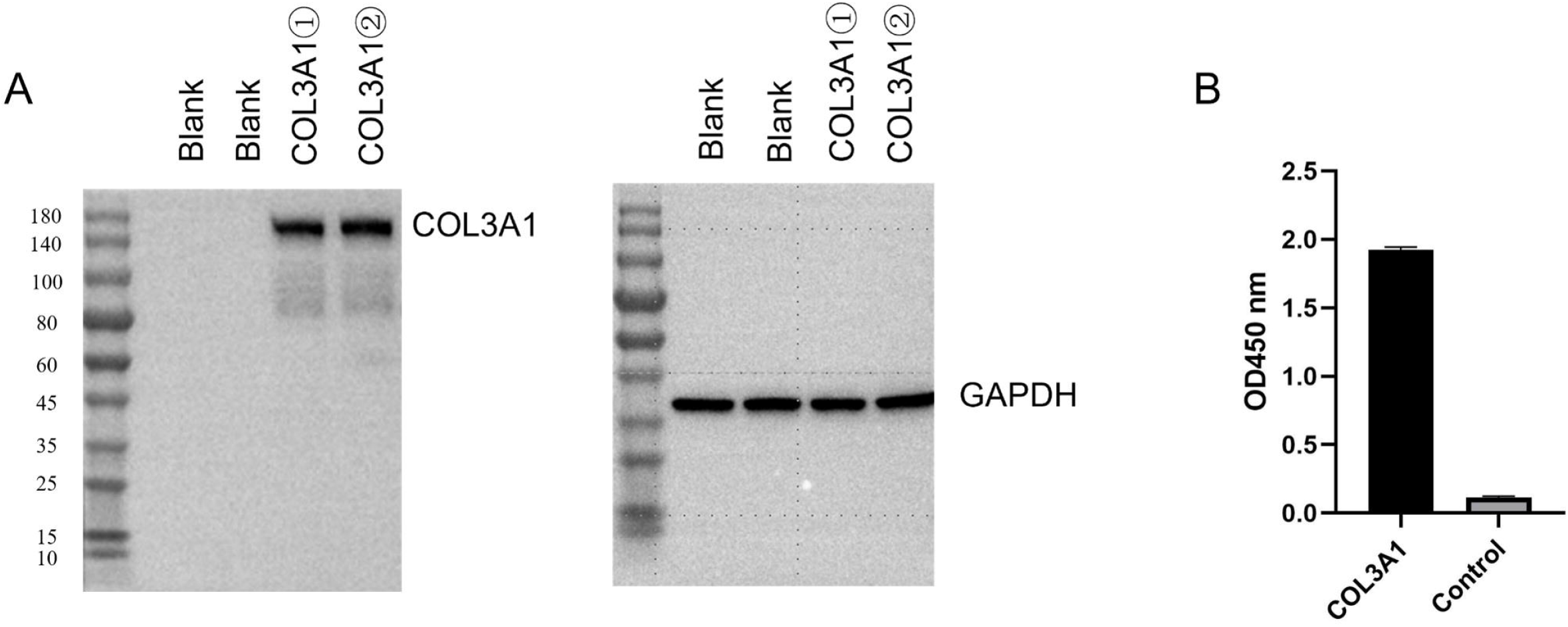
The express verification of collagen mRNA. A. Collagen type III expression detection by western blot. B. Collagen type III expression detection by ELISA.

### 2.2 *in vivo* Expression of topically applied collagen mRNA

It is critical for cosmetic applications that mRNA encoding collagens can be transdermal delivered and functionally translated. To demonstrate the sequences-optimized mRNA encoding collagens can be topically delivered and efficiently expressed *in vivo*, we engineered type III collagen mRNA with a C-terminal Flag tag. Compared to mock control, successful transdermal delivery of mRNA encoding type III collagen were confirmed by qPCR (Forward primer: ACGCAAGGCTGTGAGACTAC; Reverse Primer: CCTTTGTCATCATCGTCCTTGT) (Fig. 3A). Immunohistochemistry staining revealed Flag-positive signals, with protein signal intensity positively correlating with application duration (Fig. 3B), which confirmed *in vivo* translation and sustained expression of topically delivered mRNA encoding type III collagen (Fig. 3B).

**Figure 3.**
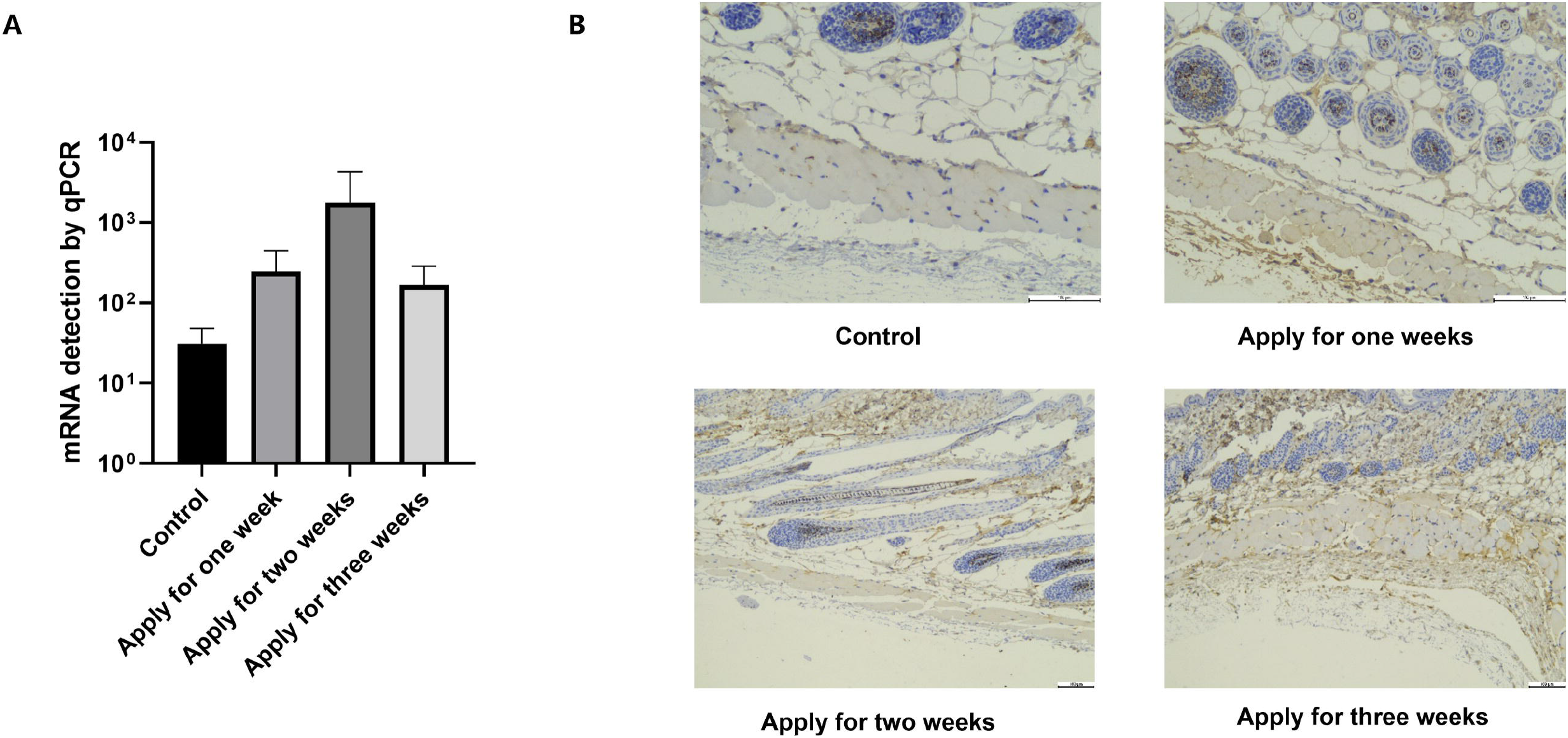
Collagen mRNA transdermal and expression verification. A. Transdermal validation of nucleic acid levels by qPCR. B. Express verification by immunohistochemical staining.

### 2.3 *in situ* Assembly of expressed collagens

Hydroxyproline, a collagen-specific amino acid, is a hallmark of collagen expression and high-order assembly. Topical application of types I or III collagen mRNA significantly increased hydroxyproline levels in skin (Fig. 4), indicating enhanced collagen synthesis and assemble. These results confirm that mRNA delivery promotes functional collagen expression in vivo, supporting skin improvement.

**Figure 4.**
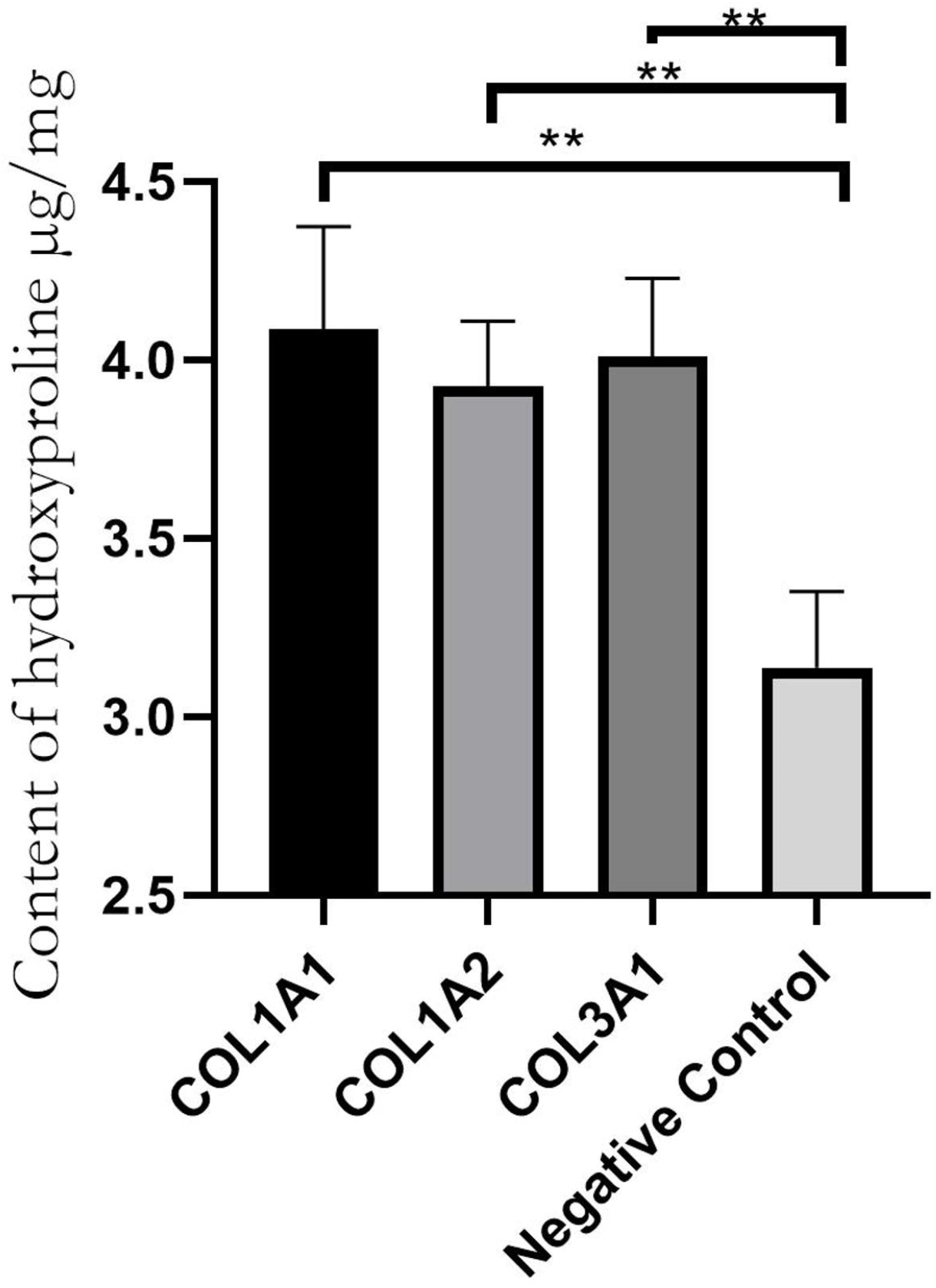
Hydroxyproline content determination. Data represent mean ± SD from three biological replicates (Student’s t test, *P < 0.05, **P < 0.01, ***P < 0.001).

### 2.4 Serum Formulation Screening

Cosmetic application of mRNA encoding collagens needs serum formulation with optimal solvents balancing irritation, transdermal efficiency, and mRNA stability. We tested ingredients combinations in serum formulations with different polypeptides and polyols. After screening serum formulations through skin sensation assessment and stability assays, optimal formulation containing ingredients that can promote transdermal delivery and reserve mRNA stability were identified. Animal studies showed enhanced delivery and *in situ* expression of collagen mRNA in serum formulation (Fig. 5). We tested the potential effects of the serum formulation containing collagen mRNA in human subjects for improving facial skin appearance. The results shown that compared with the microcrystal loading of hyaluronic acid, serum containing 5 μg collagen mRNA applied to facial skin have a better effects in wrinkle repair and skin improvement (Supplement Fig 3). Additionally, assessment of the indicators of elasticity, compactness and area of wrinkles after 14 days and 21 days of skin application, indicated that there is a significant improvement in facial skin after continuous use (Table S1).

**Figure 5.**
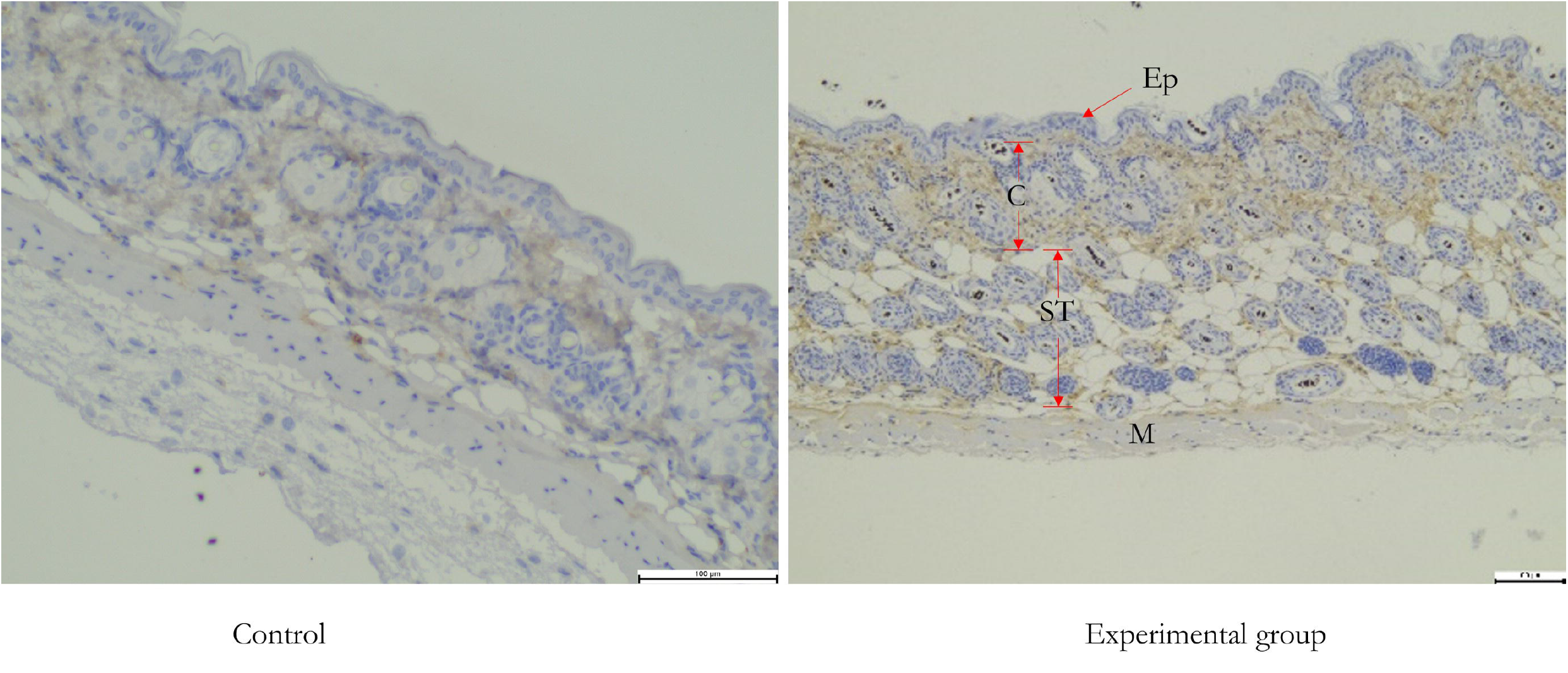
Formulation screening and express verification. EP: epidermis, C: corium layer, ST: subcutaneous tissue, M: muscular layer

## 3 Discussion

Collagen in skin can be damaged by multiple factors such as ultraviolet irradiation, mechanical stress, inflammation, and aging. Bad habits like smoking, excessive alcohol consumption, and poor diet contribute to collagen loss, which is also a natural part of aging that slows down the body’s ability to create new collagen.

Therefore, collagen supplements were regarded as important approach to improve skin moisture, elasticity and strength. Traditionally, cosmetic collagen supplements have been derived from animal sources like cows and pigs, primarily using connective tissues such as hide, skin and bones. Animal collagen is often broken down into hydrolyzed collagen (peptides) for topical products. Such approaches posed the risk of transmitting diseases such as mad cow disease and foot-and-mouth disease. At present, marine-derived collagen is considered relatively safe and an alternative source for cosmetic collagen. However, supplement collagens lack high-order structure and have poor skin-penetrating potential[14].

Topically delivering mRNA encoding collagens to the skin is a potential approach to enhance natural collagen production. The effects of rational designing of optimal secondary structure of nucleic acid to enhance delivery through intact human skin were reported by several groups and have been successfully applied to development of functional nucleic acid-based drugs[15, 16]. In this study, we demonstrated that an optimized serum formulation enables efficient transdermal delivery of in vitro-synthesized, AI-designed collagen mRNA. The delivered mRNA is functionally expressed in vivo, leading to the in situ assembly of collagens. Our works provide a practical approach for cosmetic application of mRNA technology for endogenous production of collagens in natural conformation.

## 4 Materials and Methods

### mRNA Synthesis

mRNA encoding human collagens (including type I, III and XVII) were synthesized by *in vitro* transcription (IVT). Briefly, the mRNA was produced by T7-polymerase-based *in vitro* run-off transcription.

### HDF-α cell culture

Human fibroblast cell line HDF-α were purchased from OriCell (Catalogue number HXXFB-00001). The cells were cultured with Opti-MEM medium (Thermo Fisher Scientific, Cat. 31985062), supplemented with 10% FBS, pen-strep in 37□ incubators with 5% CO_2_, as protocol provided by supplier.

### mRNA transfection and preparation of lysate

The fibroblast cell HDF-α in the logarithmic phase of growth were harvested by digesting with Trypsin containing 0.25% EDTA. Harvested cells were plated in 6-well plate with 5×10^5^ cells/well and incubated overnight in 37□ incubator with 5% CO_2_. Lipofectamine™ Messenger MAX™ (lipo MAX) (Invitorgen, Cat. LMRNA008) were used for mRNA transfection as protocol provided by supplier. Briefly, the transfection complex was prepared by mixing 3.75μl lipo MAX in 15μl Opti-MEM with 2μg mRNA in 12μl Opti-MEM, then added to cells (250μl/well) in 6-well plate, incubating in a 37□ incubator overnight. Cells were harvested, washed with PBS and then lysed with Cell lysis buffer (Beyotime, Cat. P0013). The Pierce™CA Protein Assay Kit was used to measure the total protein concentration and normalize it to 1 mg/ml. To quantify the potential of optimized mRNA to improve collagen expression in fibroblast cells, mRNA containing serum was directly mixed with HDF-α cells following the protocol of Group Standard T/SHRH 031-2020 issued by Shanghai Daily Chemistry Trade Association[12].

### Western blot

Protein concentration was measured by BCA method, equal amount of protein samples was used, 5×Loading Buffer was added, and the samples were denatured by boiling at 95□ for 10 min, then centrifuged. The separating gel and the stacking gel were prepared and combed, and the coagulation time was required to be >2 hours. 15-25μL of samples and pre-stained Marker were added to each well, and electrophoresis was performed until the interface of the concentration gel and the stacking gel, then the voltage was adjusted to 100-120V, until the targets migrated to 2/3 of the separating gel. The PVDF membrane was soaked in methanol for 1 min, and then soaked in transfer buffer with gel and filter paper for 15 min. The transfer clamp was assembled in order, and the transfer was carried out for 40 minutes. 5% skim milk was incubated at room temperature for 2 hours. The primary antibody (abcam, ab308455; Cell Signaling technology 30565S; Abclonal A25310 and A4808) was diluted according to the instructions, added and incubated at room temperature for 2 hours, then the primary antibody was recovered; the secondary antibody (Proteintech, SA00001-2) was diluted 1:2000, and incubated at room temperature for 1 hour. The ECL A/B solution was mixed, covered the surface of the membrane, and by fluorescence scanner.

### qPCR

cDNA, specific primers, fluorescent dyes, dNTPs, MgCl□, Taq DNA polymerase and other enzymes and buffers are mixed. A thermal cycling reaction is performed: denaturation at 95□, annealing at 58□, and DNA chain extension catalyzed by Taq enzyme at 72□, during which the dyes produce a fluorescent signal. The fluorescence intensity is detected during the extension phase of each cycle, and the data is recorded and analyzed in real time The cycle number (Ct value) at which the sample reaches the threshold is read, and the results are calculated and compared with the control. The specific primers (COL3A1-3XF-F3: ACGCAAGGCTGTGAGACTAC, COL3A1-3XF-R3-2: CCTTTGTCATCATCGTCCTTGT) were employed in nucleic acid identification.

### Immunohistochemical staining

Fix the tissue section samples, use blocking agents to reduce the possibility of non-specific binding, and ensure that the antibody only binds to the target ant. Add antibodies conjugated with chromogenic agents (SinoBiological, 109143-MM13-H) specific to certain antigens, allowing them to bind to the antigens in the tissue sections. Remove unbound antibodies to reduce background staining. Visualize the location of the antigen through a chemical reaction that causes the chromogenic agent to be visible. For better observation of cellular, counterstaining can be performed, and the sections can be mounted for microscopic examination.

### Elisa

ELISA detection of COL3A1: Coated with 2ug/ml of flag antibody (abcam, ab125243), 100ul/well, 4 ℃ overnight; after washing the plate, add 100μl/well of blocking solution, incubate at 37□ for 2h; after washing the plate, add diluted samples, 100ul/well, incubate at 37□ for 2h; after washing the plate, add diluted COL3A1 antibody (CellSignalingTechnology, 30565), 100μl/well, incubate 37□ for 1h; after washing the plate, add diluted HRP-anti rabbit IgG antibody (abcam, ab97051), 100μl/well, incubate at 37□ for 1h; after washing the plate, add color developing solution to develop color for 10min, and then measure the OD value at 450 with an enzyme-linked immunosorbent assay after termination.

### Animal Experiments

Animal experiments were approved by Institutional Review Board of Rhegen. For topical application of mRNA encoding collogens, BALB/C mice were shaved and further applied with the depilatory cream for topical delivery experiment of mRNA containing serum. After anesthesia, mRNA containing serum were evenly applied to the depilated area followed by gentle rubbing with a clean pad.

### Skin condition improvement study by volunteers

The study to assess skin condition improvement in volunteers was conducted after obtaining approval from the Institutional Review Board of Rhegen. After cleansing, one bottle of the serum preparation was evenly applied to the skin around the eyes, followed by gently massage to assist absorption of the lipid. Volunteers used the serum containing mRNA twice a day with each in the morning and the evening for 21 consecutive days. Control samples were applied to volunteers with moisturizing serum using a microcrystal infusion instrument. The measured values of each test area were then statistically analyzed.

## Supporting information

The express verification of collagen type I and type XVII mRNA

in vitro evaluation of transdermal delivery of mRNA encoding collagen type I A. relative expression of collagen I of mRNA with direct incubate with fi

Product efficacy verification in human body. A. Product effect display after continuous application 5 ug collagen mRNA. B. Product effect display afte

Skin improvement indicators statistics.

---

**Supplement Figure 1 The express verification of collagen type I and type XVII mRNA**

**Supplement Figure 2 *in vitro* evaluation of transdermal delivery of mRNA encoding collagen type I** A. relative expression of collagen I of mRNA with direct incubate with fibroblasts according to Group Standard T/SHRH 031-2020, B. expression of collagen I of mRNA in T-Skin™ model.

**Supplement Figure 3 Product efficacy verification in human body**. A. Product effect display after continuous application 5 ug collagen mRNA. B. Product effect display after microcrystalline imported hyaluronic acid essence.

**Supplement table 1 Skin improvement indicators statistics**.

