## Supplementary figures and images for "Delivery of collagen-encoding mRNA to the skin for cosmetic applications"

### in vitro evaluation of transdermal delivery of mRNA encoding collagen type I A. relative expression of collagen I of mRNA with direct incubate with fi

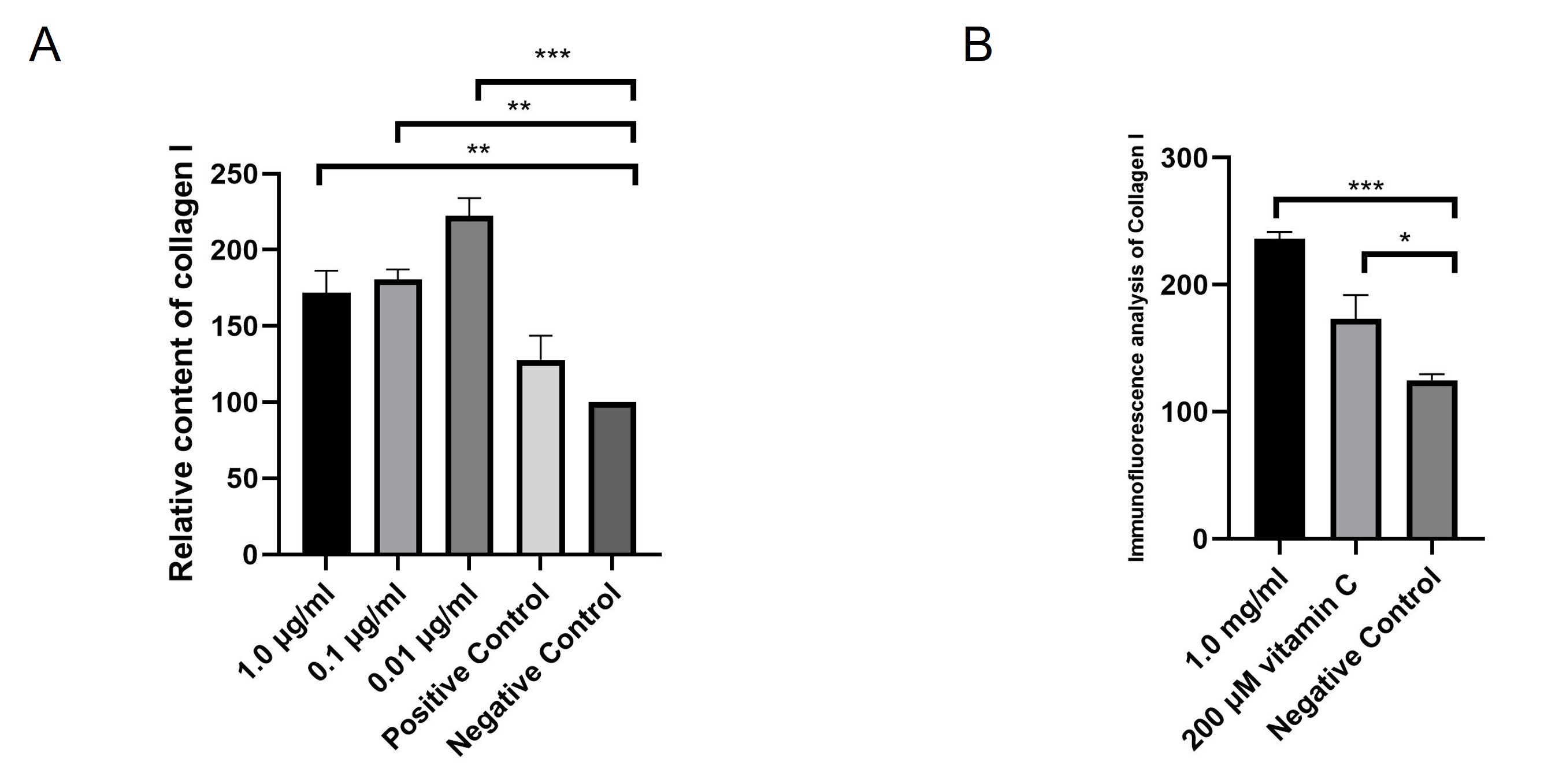

### Product efficacy verification in human body. A. Product effect display after continuous application 5 ug collagen mRNA. B. Product effect display afte

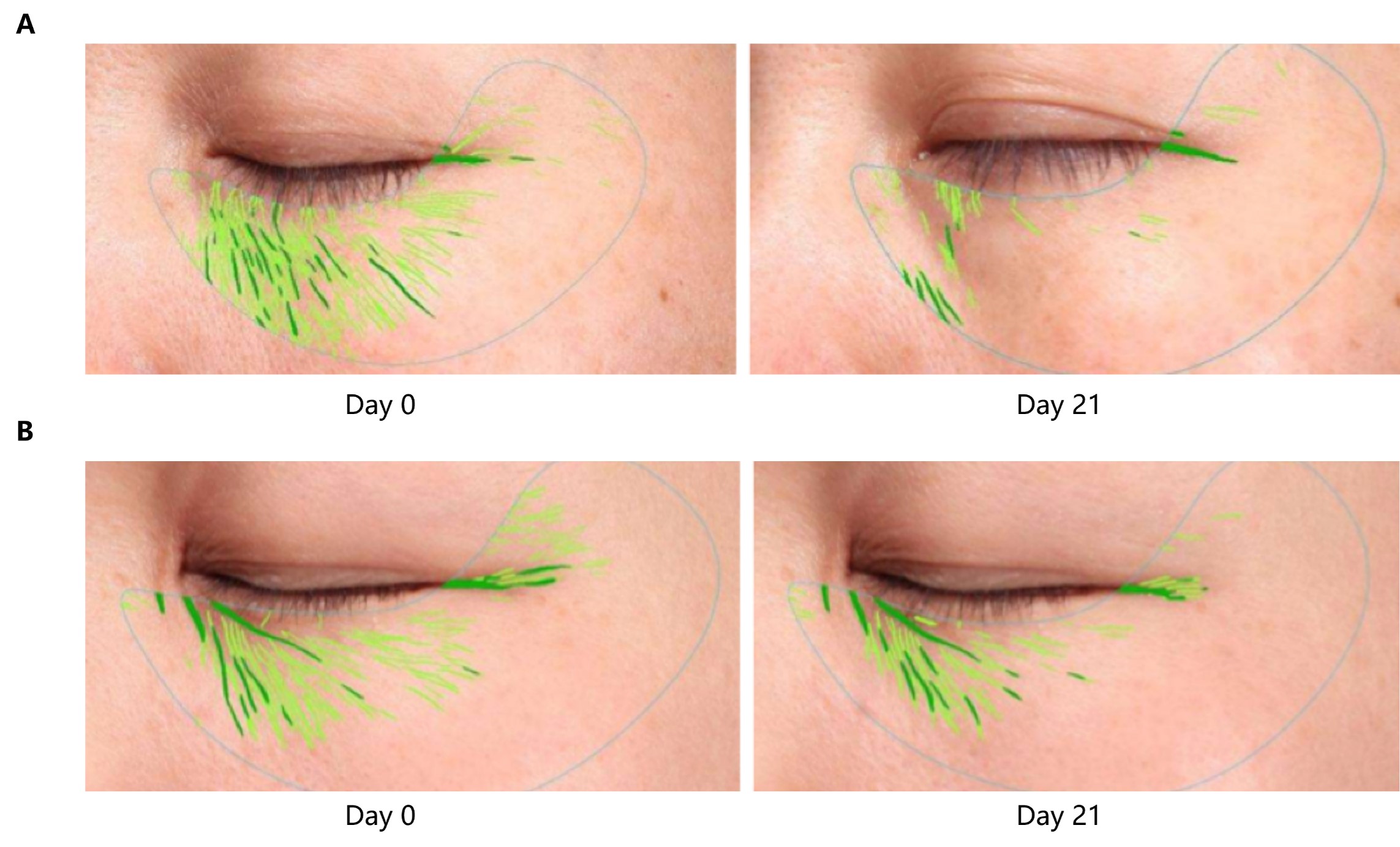

### The express verification of collagen type I and type XVII mRNA

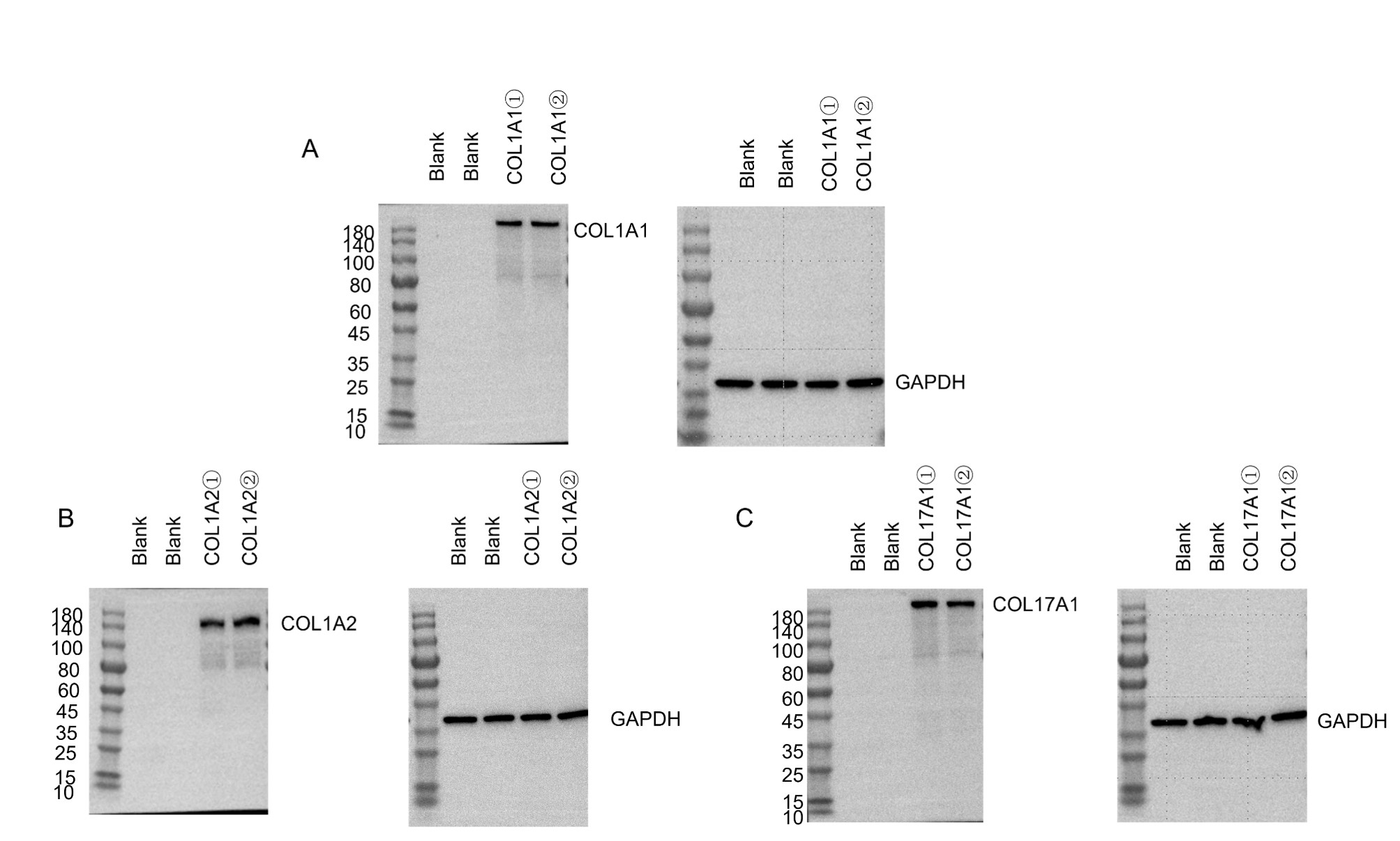
